# Towards Reliable Tracking of Natural Killer Cells Using Commercial Iron Oxide Nanoparticles and Magnetic Particle Imaging

**DOI:** 10.64898/2026.04.24.720698

**Authors:** Suzanne Lightsey, Victoria Consalvo, SK Rajab Ali, Daniela Paula Valdés, Jeremiah Oyer, Grace Gloger, Alicja Copik, Carlos Rinaldi-Ramos, Blanka Sharma

## Abstract

Non-invasive tracking of natural killer (NK) cells remains a major challenge in cancer immunotherapy, limiting our understanding of their in vivo migration and persistence. Magnetic particle imaging (MPI) offers a quantitative, real-time method for visualizing labeled cells, yet optimal labeling protocols for NK cells have not been established. Here, we evaluate commercially available iron oxide nanoparticles (IONPs) for MPI labeling of both NK92MI cells and primary human NK cells. Labeled cells retained viability and cytotoxicity, including activity against three-dimensional tumor spheroids, and were detectable by MPI. To further examine imaging performance in a biologically relevant context, we employed mouse phantoms that recapitulate organ-specific signal distributions, enabling evaluation of quantification and liver spillover effects. We identify key tradeoffs between particle colloidal stability and per-cell iron content: VivoTrax and VivoTrax Plus provided higher MPI signal but required post-labeling purification, reducing cell recovery, whereas Synomag-D and Perimag were more stable and preserved cell yield despite lower signal intensity per cell. These results provide a framework for selecting nanoparticles that balance detection sensitivity, cell viability, and workflow practicality, advancing non-invasive NK cell tracking.

## Introduction

Adoptive cell-based immunotherapies have been transformative in cancer treatment; however, their success remains variable across cancer types. While T cell–based adoptive cell therapies (ACT) have led to remarkable successes in certain hematologic malignancies, their impact in solid tumors has been limited by poor infiltration, antigen escape, and safety risks such as graft-versus-host disease[1–3]. Natural killer (NK) cells offer a promising alternative, as they can target tumor cells without prior antigen sensitization or MHC restriction and carry a lower risk of severe immune-related toxicities, making them well-suited for off-the-shelf, allogenic use[4–7]. However, despite these advantages, NK cell– based therapies face their own barriers, including limited total in vivo persistence and poor accumulation in solid tumors[8]. Understanding how NK cells traffic, localize, and function within tumor sites remains a major challenge in the field[9,10]. As a result, there is growing interest in developing non-invasive, quantitative methods to track NK cells in vivo, enabling researchers to evaluate their biodistribution and infiltration over time[11]. Such tracking technologies are critical not only for understanding the behavior of NK cells in complex tumor microenvironments but also for guiding the rational design of next-generation NK cell therapies with improved efficacy in solid tumors.

Several imaging platforms have been explored for tracking the biodistribution and fate of adoptively transferred cells in vivo, including fluorescence and bioluminescence imaging, magnetic resonance imaging (MRI), positron emission tomography (PET), and single-photon emission computed tomography (SPECT). These techniques rely on either reporter gene expression to enable signal generation within living cells[12] or the use of externally applied labeling agents to provide detectable contrast[13]. For instance, iron oxide particles are often used for MRI-based cell tracking[14,15], while PET and SPECT utilize radiolabeled tracers to monitor cellular localization[16]. Despite their widespread application, many of these methods are hindered by challenges such as poor spatial resolution, limited tissue penetration, and signal ambiguity caused by background or artifacts. Furthermore, the tracers used are often limited by poor stability and dose-limiting toxicities, which hinder both sensitivity and the observation window[12,17]. These constraints can obscure interpretation, particularly when tracking cells in deep tissues or complex tumor microenvironments[12,18].

Among emerging technologies, magnetic particle imaging (MPI) offers several advantages that make it particularly well-suited for tracking cell-based therapies. First introduced in 2005[19], MPI enables the direct detection of iron oxide nanoparticles (IONPs), producing positive contrast images that are inherently free from background tissue signal. Unlike MRI, which relies on indirect contrast mechanisms and is prone to signal voids and susceptibility artifacts, MPI generates signal that scales linearly with the amount of IONP present, facilitating accurate and quantitative assessments of labeled cells[12,20]. In addition to its quantification capabilities, MPI is not limited by depth attenuation and does not rely on ionizing radiation, making it particularly well-suited for longitudinal, in vivo studies[19]. IONPs used for MPI are generally biocompatible and, once internalized by cells, are naturally processed through physiological iron metabolism pathways, with iron stored in hemoglobin or ferritin[21,22]. Moreover, MPI data can be co-registered with anatomical imaging modalities such as computed tomography (CT) or MRI, enabling integration of functional and structural information. With several human-scale MPI systems currently under development[23–27], the technology is progressing rapidly toward clinical translation, offering a compelling platform for noninvasive, quantitative monitoring of adoptive immune cell therapies.

MPI has been successfully applied to track various immune cell types, including dendritic cells[28], macrophages[29], and T cells[12,30]. However, labeling NK cells for MPI presents unique challenges due to their low endocytic activity and limited cytoplasmic volume, both of which restrict efficient uptake of nanoparticles. In a study by Sehl et al., the feasibility of labeling NK92 cells, a human NK cell line, for MPI was demonstrated using VivoTrax, a commercially available IONP formulation, where the cells achieved an average uptake of 3.19 picograms of iron per cell[31]. While this confirmed the potential for IONP-based labeling of NK cells, a substantial fraction of the MPI signal originated from extracellular IONP aggregates, producing signal unrelated to the cells themselves[31]. Since NK cells are non-adherent and remain in suspension culture, it is particularly difficult to separate free tracer from intracellular labeled cells during post-labeling washes. As a result, residual extracellular IONP can confound in vivo imaging by generating signal that does not accurately reflect the location or quantity of labeled NK cells.

In this study, we evaluated four commercially available IONPs under various culture conditions to identify protocols that promote IONP uptake, preserve NK cell viability and cytolytic function, and produce reliable MPI signal. We found that the choice of culture media had minimal impact on IONP stability and NK cell labeling efficiency. Among the particles tested, Perimag and Synomag-D demonstrated high colloidal stability but yielded relatively low MPI signal per cell, requiring higher cell numbers for effective imaging. In contrast, VivoTrax and VivoTrax Plus generated stronger MPI signals but were prone to aggregation, which complicated interpretation and could introduce non-cell-associated signal. To address this, density gradient separation was used to reduce extracellular iron in VivoTrax Plus-labeled samples. Our findings were consistent across both NK92MI cells, an IL2-independent NK cell line, and ex vivo expanded primary NK cells. Furthermore, we assessed the imaging capabilities of IONP-labeled NK cells using MPI in a biologically relevant setting with an anatomically accurate mouse phantom. Together, this work establishes key considerations for optimizing IONP labeling of NK cells for MPI and represents a step toward enabling accurate, non-invasive tracking of NK cell therapies in vivo.

## Materials and Methods

### IONP Characterization and Stability

Commercial IONPs Perimag plain (Micromod, 102-00-132), Synomag-D plain 50 nm (Micromod, 104-00-501), VivoTrax (Magnetic Insight), and VivoTrax Plus (Magnetic Insight) were obtained from their suppliers and stored at 4°C. IONPs were imaged using a FEI Talos F200i S/TEM instrument to evaluate particle size and morphology. 20 μL of IONPs were loaded onto a 200-mesh carbon film with copper grids and allowed to dry. Grids were imaged at 200 kV and up to 300 kx magnification. The colloidal stability of commercial IONPs in NK cell media (RPMI 1640 + 1% v/v penicillin-streptomycin, RPMI 1640 + 10% v/v FBS + 1% v/v penicillin-streptomycin, and OptiMEM + 1% v/v insulin transferrin selenium (ITS) + 1% v/v penicillin-streptomycin) was measured using dynamic light scattering (DLS) measurements in a Wyatt DynaPro over 70 hours at room temperature (n=3).

Iron concentrations of the IONPs were quantified using a colorimetric assay based on 1,10-phenanthroline complexation, as previously described[20]. Samples were digested in nitric acid, then treated with hydroxylamine hydrochloride to reduce Fe^3+^ to Fe^2+^, followed by sodium acetate and 1,10-phenanthroline to form the colored complex. Absorbance was measured at 508 nm, and concentrations were calculated using a standard calibration curve. All measurements were performed in triplicate.

### Primary NK Cell Isolation and Expansion

PM21-expanded primary NK cells were kindly provided by the Copik Laboratory (University of Central Florida, Orlando, FL). These cells were generated from peripheral blood mononuclear cells (PBMCs) using plasma membrane particles (PM21) derived from K562 cells expressing membrane-bound IL21 and 4-1BBL, as previously described[32,33]. Briefly, PBMCs from de-identified healthy donors were expanded over 14 days in SCGM supplemented with IL2, IL12, IL15, IL18, and PM21 particles. Media and cytokine conditions were refreshed every 2-3 days, and cells were transitioned to RPMI 1640 media during the second week of culture. The preparation and characterization of PM21 particles, as well as the full expansion protocol, followed established methods outlined in prior studies[32,33].

### Cell Culture

NK92MI (ATCC, CRL-2408) and PM21 NK cells were maintained at 1×10^6^ cells/mL in RPMI 1640 supplemented with 10% (v/v) fetal bovine serum (FBS)(Corning) and 1% (v/v) penicillin-streptomycin (Gibco) at 37°C in a humidified incubator containing 5% CO_2_. Recombinant human IL2 (Peprotech, 100U/mL) was added daily to primary NK cell culture.

### Cell Labeling

NK cells at 1.2×10^6^ cells/mL were incubated with 100 μg Fe/mL of commercial IONPs for 24 hours at 37°C in a humidified atmosphere containing 5% CO_2_. Labeling was performed in three different culture media: (1) RPMI 1640 supplemented with 1% (v/v) penicillin-streptomycin, (2) RPMI 1640 supplemented with 10% (v/v) FBS and 1% (v/v) penicillin-streptomycin, and (3) OptiMEM supplemented with 1% (v/v) ITS and 1% (v/v) penicillin-streptomycin[31,34,35]. Following incubation, cells were washed three times with phosphate-buffered saline (PBS), gently resuspended, and centrifuged at 450xg for 3 minutes between washes to remove extracellular nanoparticles.

### Cell Viability

IONP labeled-NK92MI cells were stained with the LIVE/DEAD™ Fixable Near-IR Dead Cell Stain Kit (ThermoFisher Scientific) according to manufacturer protocols. A minimum of 10,000 cells were analyzed using a BD FACS Celesta Flow Cytometer and Flowjo Software v10 (FLOWJO, LLC Data analysis software). PM21 NK cell viability after incubation with IONPs was determined using either trypan blue exclusion assays or proliferation assays, as previously described[34]. Briefly, PM21 NK cells were labeled with Cell Trace Violet (ThermoFisher Scientific) and proliferation was assessed using a CytoFlex LX flow cytometer to determine the cells’ doubling time.

### Functional Assays

The cytotoxic activity of PM21-expanded NK cells from two donors against K562-GFPLuc target cells was assessed using an Annexin V assay, as previously described[34]. Briefly, NK cells were incubated with 100 μg Fe/mL of IONPs for 24 hours in RPMI 1640, supplemented with 10% FBS and 1% penicillin-streptomycin, prior to the assay. NK cells were co-cultured with K562-GFPLuc target cells at various effector to target (E:T) ratios for 60 minutes at 37°C in a humidified tissue culture incubator with 5% CO_2_. Following incubation, cells were harvested by centrifugation and stained with Annexin V-Pacific Blue (BioLegend) for 15 minutes at 4°C in the dark, according to the manufacturer’s protocol. Samples were subsequently analyzed by flow cytometry and cytotoxicity was assessed by quantifying the absolute number of viable target cells (VTC), defined as GFP^+^/Annexin V^-^, in each well containing NK cells (VTC^E:T^). Cytotoxicity was calculated relative to the average number of viable targets in wells containing target cells alone (VTC^T^), using the following equation:

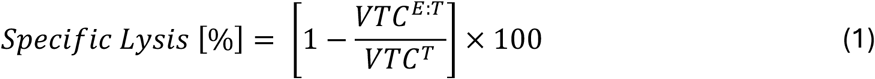

Kinetic cytotoxicity assays were performed using the IncuCyte Live-Cell Analysis System (Sartorius), as previously described[36]. Cancer cell lines expressing green fluorescent protein (GFP), including A549-GFP and NCI-H1299-GFP, were seeded in ultra-low attachment plates to form 3D spheroids. PM21 NK cells were labeled with 100 μg Fe/mL of IONPs for 24 hours in RPMI 1640 supplemented with 10% (v/v) FBS and 1% (v/v) penicillin-streptomycin. Following labeling, NK cells were co-cultured with tumor spheroids at varying E:T ratios for up to 7 days. Tumor growth was monitored by measuring green object integrated intensity (GOII). To evaluate cytotoxicity, GOII at each time point (GOII_t_) was normalized to baseline values at the time of NK cell addition (GOII_t=0_), yielding normalized GOII (nGOII = GOII_t_ / GOII_t=0_). Cytotoxicity (%) was calculated using the following equation:

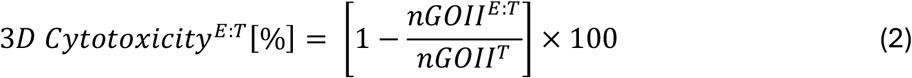

EC_50_ (half-maximal effective concentration) values were used to compare NK cell cytotoxicity across conditions, defined as the NK cell concentration required to achieve a response halfway between baseline and maximal killing.

### Density Gradient Separation

Following a 24-hour incubation of NK cells with 100 μg Fe/mL VivoTrax Plus IONPs in RPMI 1640 supplemented with 10% (v/v) FBS and 1% (v/v) penicillin-streptomycin, the cell suspension was carefully layered over 3 mL of Ficoll-Paque Premium (1.078 g/mL, Cytiva). Samples were centrifuged at 450xg for 20 minutes at room temperature without brakes to separate free iron aggregates from the labeled cells. NK cells were collected from the interface between the Ficoll and the overlaid medium[37].

### Microscopy

IONP-labeled NK cells were fixed in methanol for 20 minutes at room temperature. After fixation, cells were washed twice with ultrapure water, centrifuging at 450xg for 3 minutes between washes. For Prussian blue staining, cells were first incubated in 14% hydrochloric acid for 10 minutes, after which an equal volume of 2.5% potassium ferrocyanide (Abcam) was added, and samples were incubated for an additional 20 minutes. Cells were then washed twice with ultrapure water and counterstained with Nuclear Fast Red (Abcam) for 10 minutes. A final series of three washes with ultrapure water was performed, centrifuging at 450xg for 3 minutes each. Stained cells were transferred to a glass-bottom well plate and imaged using a Keyence BZ-X710 epifluorescence microscope.

### Magnetic Particle Imaging and Processing

To assess MPI signal, samples were transferred to 0.6 mL microcentrifuge tubes and imaged using a MOMENTUM scanner (Magnetic Insight) in high-sensitivity mode. Two-dimensional scans were performed at 45 kHz with 15.5/20.5 mT excitation in the x/z axes and a gradient strength of 3.055 T/m. DICOM files obtained through X-space direct reconstruction and the application of an equalization filter[38] were analyzed by searching for the maximum signal s_max_ in the scan and thresholding up to 0.5s_max_ (0.5Max threshold), defining regions of interest (ROIs). Peak signal-to-noise ratio (pSNR) was calculated by dividing the maximum signal intensity within a 15 mm circular ROI by the standard deviation of an empty (background) scan. Signals with a pSNR greater than 3 were considered detectable. Total MPI signal (S) was calculated by multiplying the average voxel intensity within each ROI ⟨*s*⟩ by the total number of voxels (N) in the segment, S=N⟨*s*⟩. Calibration curves were generated by plotting total MPI signal against known iron mass from reference samples (Supplemental Figure S1). A linear regression was applied to derive an equation relating total signal to iron mass, which was subsequently used to estimate IONP content in experimental samples. Triplicates of all samples were prepared and measured; the informed results are the average these measurements.

For evaluation of IONP-labeled cells under biomedically relevant conditions, a 3D-printed anatomically correct mouse phantom was used[39,40]. The phantoms included a fillable liver cavity and ports in the brain and lung regions. Briefly, the mouse phantom was printed with the Form 3 stereolithography (SLA) printer (Formlabs Inc.) using Clear V4.0 resin (Formlabs Inc.) at a layer thickness of 50 μm. Following printing, the phantoms were washed with isopropyl alcohol (IPA), with additional flushing of the liver cavity and ports in the brain and lung regions to ensure removal of residual resin. The models were then dried with compressed air and post-cured in the Form Cure (Formlabs Inc.) using a 405 nm light at 60°C for 25 minutes. For each phantom, 4.5×10^6^ NK cells labeled with either Synomag-D or separated VivoTrax Plus IONPs were injected into the liver cavity. 2.25×10^5^ of IONP-labeled NK cells were placed in a capillary inserted into either the brain or lung port. Cell numbers were kept constant across conditions to directly compare the performance of the tracers. Phantoms were scanned in 2D in the MOMENTUM scanner in high-sensitivity mode (45 kHz with 15.5/20.5 mT excitation amplitudes in the x/z axes and a gradient strength of 3.055 T/m). The signals in the brain and lung regions were thresholded with the 0.5Max criterion, and the total signal was calculated again as S=N⟨*s*⟩. To evaluate the influence of spillover signal from the liver, a circle centered at the local maximum with a fixed diameter of 7 mm was placed in the position corresponding to the maximum signal intensity in the ROI.

### Statistical Analysis

Statistical analysis was performed by GraphPad Prism 10.6.1. Unpaired two-tailed Student’s t-tests were used unless noted in the figure legend. All experiments were performed for at least 3 biological replicates. Error bars represent the standard deviation of the mean. Statistical comparisons were considered significant if p < 0.05. P values are shown as * if p< 0.05, ** if p < 0.01, *** if p < 0.001, and **** if p < 0.0001.

### Data Availability

All data supporting the results in this study are available within the article or included in the Supplementary Materials. Access to raw data sets is available from the corresponding author upon reasonable request.

## Results

### Commercial IONP Characterization and Stability

To evaluate the suitability of IONPs for NK cell labeling and MPI detection, four common, commercially available nanoparticles were compared: Perimag, Synomag-D, VivoTrax, and VivoTrax Plus. These particles differ in core structure and surface coating, with Perimag consisting of clustered dextran-coated iron oxide, Synomag-D composed of maghemite nanoflowers coated with dextran, and both VivoTrax formulations containing iron oxide magnetite cores coated with carboxydextran, with VivoTrax Plus representing a filtered version enriched for larger cores[20]. Representative TEM images showed distinct particle morphologies (Figure 1A), with Synomag-D displaying the expected nanoflower morphology, while Perimag, VivoTrax, and VivoTrax Plus contain smaller cores that form more amorphous and heterogeneous aggregates in both size and shape. The TEM analysis was performed to visualize nanoparticle core size and morphology, rather than to quantitatively assess larger-scale clustering. Because TEM imaging is subject to sampling bias and drying effects, the observed structures may not fully represent the morphology of particles in suspension. 2D high-sensitivity MPI scans of 20 μg Fe samples demonstrated characteristic signal patterns for each IONP (Figure 1B).

**Figure 1.**
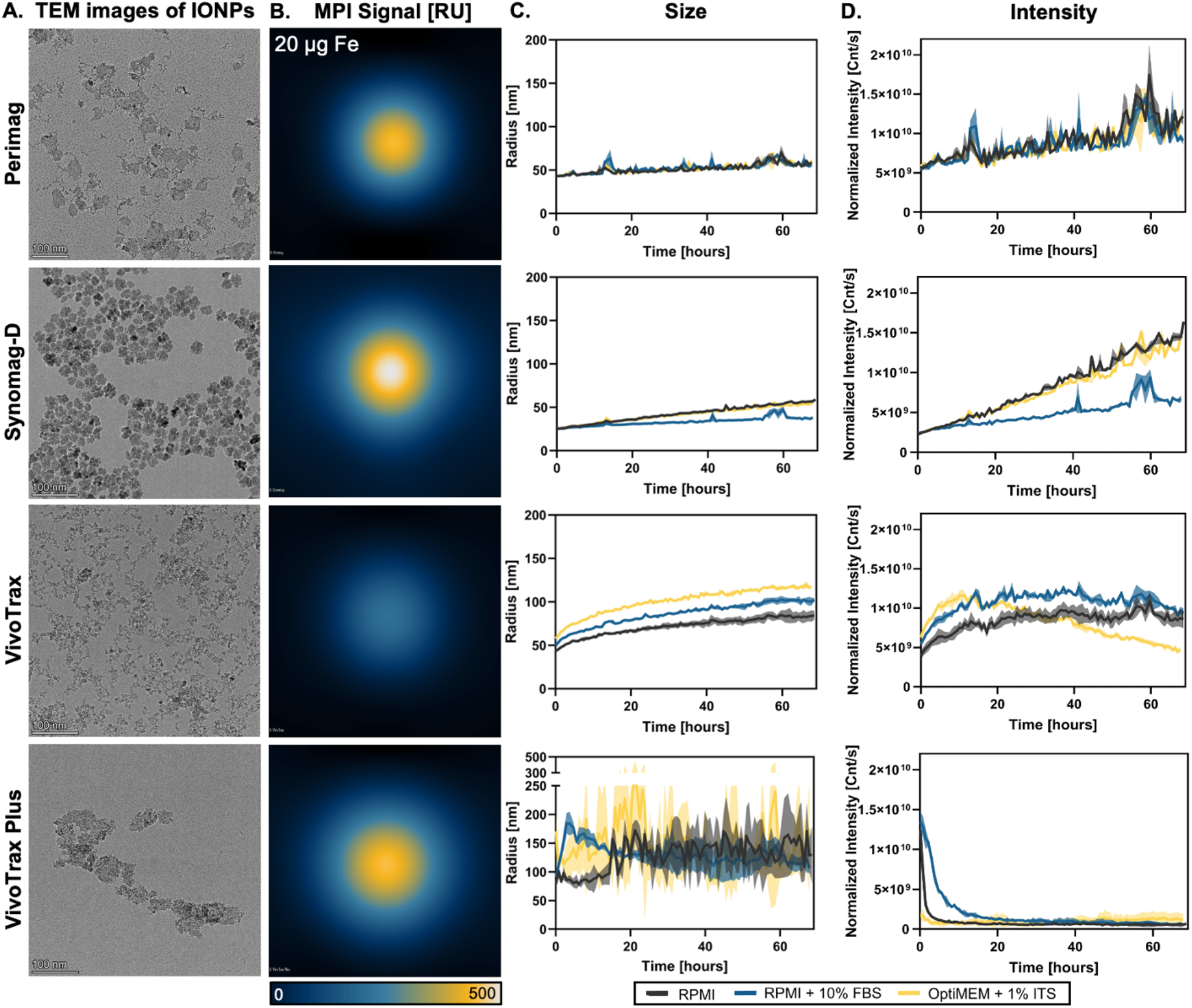
Iron Oxide Nanoparticle Characterization and Stability. (A) Representative TEM images of each IONP (scale bar = 100nm). (B) Representative 2D high-sensitivity MPI scans for samples of each IONP with 20 μg of iron. IONP stability in different NK cell culture media via (C) nanoparticle size and (D) normalized intensity over 70 hours, where an increase in size and decrease in normalized intensity indicate nanoparticle aggregation.

To assess their size and stability in relevant culture conditions, changes in the hydrodynamic radius and normalized intensity signal were monitored by DLS over 70 hours in 3 different NK cell culture mediums: RPMI 1640 (no serum), RPMI 1640 + 10% FBS (serum-containing), and OptiMEM + 1% ITS (reduced serum). Perimag and Synomag-D had relatively stable size and signal intensity across all conditions, indicating colloidal stability with minimal aggregation (Figure 1C and 1D). In contrast, both VivoTrax and VivoTrax Plus showed either a gradual increase in apparent hydrodynamic radius or substantial fluctuations in size measurements, accompanied by a marked decrease in the normalized intensity, which measures the number of photons scattered by the particles per second (Figure 1C and 1D). These trends are consistent with nanoparticle aggregation and sedimentation, where larger aggregates settle to the bottom of the well, leading to uneven particle distribution and unreliable DLS measurements over time (Supplemental Figure S2).

### In Vitro Analysis of NK92MI Cell Labeling

NK92MI cells were initially used to establish the experimental conditions necessary for iron labeling and imaging prior to transitioning to primary cells. Prussian blue staining was performed to qualitatively assess iron labeling of NK92MI cells following co-culture with 100 μg Fe/mL of each IONP in different culture media and time points (Figure 2A, Supplemental Figure S3). In VivoTrax and VivoTrax Plus conditions, extracellular iron aggregates were apparent even after multiple wash steps, suggesting incomplete removal of unbound particles. In contrast, Perimag and Synomag-D samples showed minimal visible aggregation. Cell viability remained high (>80%) across all groups, indicating that the IONPs were not cytotoxic at this concentration (Figure 2B). MPI analysis revealed low overall signal and poor signal-to-noise ratios, with detectable signal (pSNR > 3) only in samples containing visible aggregates (Figure 2C). Based on these findings, serum-containing media were selected for subsequent experiments, as this condition did not exacerbate nanoparticle aggregation and is better suited for longer-term functional studies. To further improve MPI signal-to-noise ratios, a higher number of NK cells was used in later experiments.

**Figure 2.**
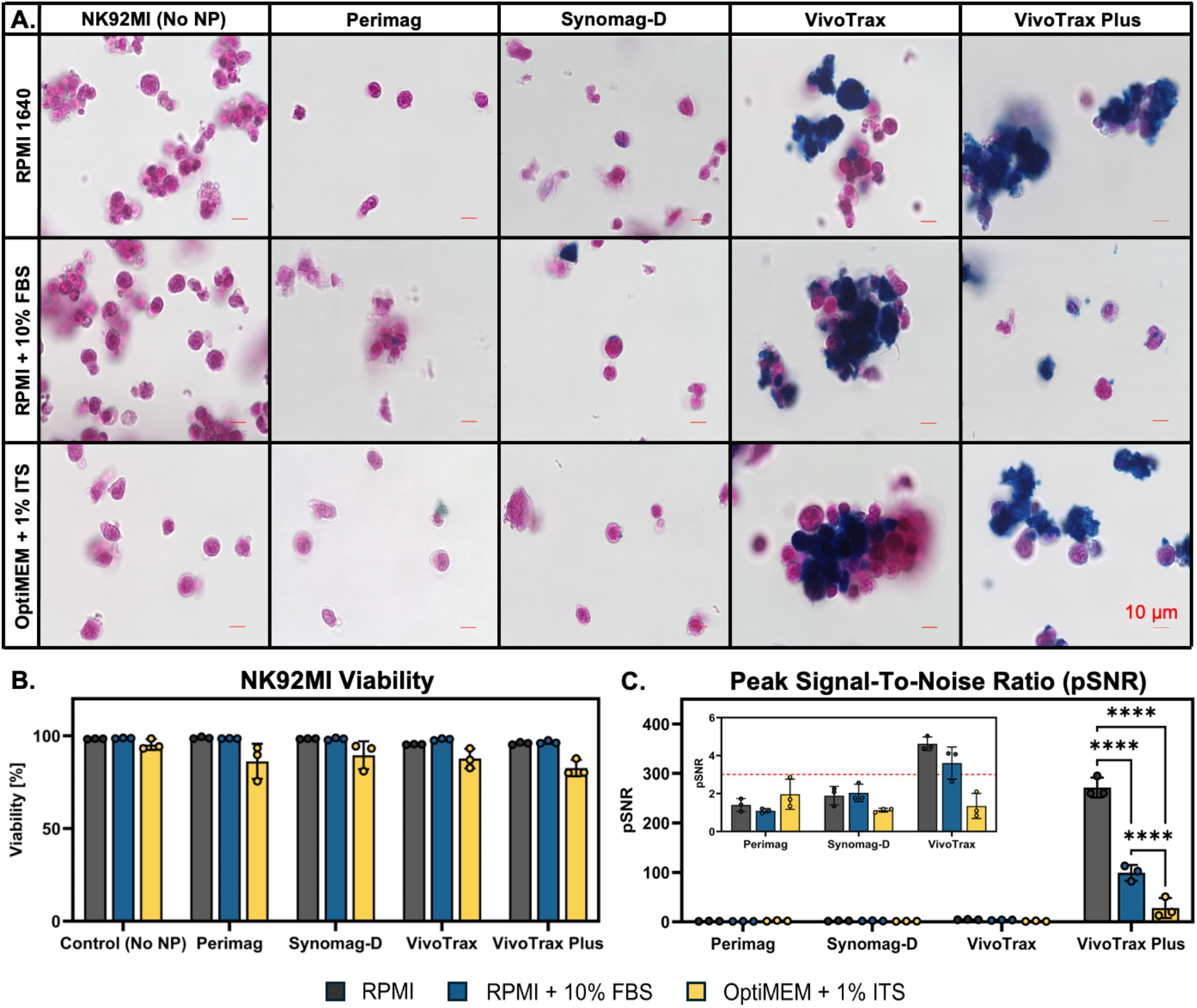
Iron Oxide Nanoparticle Interactions with NK92MI Cells. (A) Prussian blue staining shows iron (blue) associated with NK92MI (pink) after labeling with 100 μg Fe/mL IONP for 24 hours (scale bar = 10 μm) in various medias. (B) Viability [%] of NK92 MI cells after incubation with 100 μg Fe/mL of IONP in various culture medias for 24 hours at 37°C and 5% CO_2_. (C) Peak signal-to-noise (pSNR) ratio demonstrates that IONP uptake is low, and larger samples need to be used. Samples with detectable signal are defined as having a pSNR > 3, as denoted by the red dashed line. Statistical significance was determined by a two-way ANOVA followed by a Tukey’s multiple comparison test, where p < 0.0001.

### In Vitro Analysis of Primary NK Cell Labeling

In primary NK cells, uptake was assessed using cells isolated from two donors and expanded with PM21 particles and cytokines, as described by Oyer et al. (2024)[34]. Prussian blue staining confirmed that IONP interactions with primary NK cells were consistent with those observed in NK92MI cells: Perimag and Synomag-D formulations showed fewer aggregates, whereas VivoTrax and VivoTrax Plus resulted in significantly more aggregation (Figure 3A). Notably, aggregation was more pronounced in primary NK cells, potentially due to the addition of exogenous IL-2 or the smaller size of the primary cells, which complicated the washing process. Despite increased nanoparticle aggregation, primary NK cells remained viable and proliferative, and NK cells retained their cytotoxic function following IONP exposure (Figure 3B and 3C). MPI analysis revealed that colloidally stable IONP formulations produced weak signals (Perimag = 1.768 pg Fe/cell; Synomag-D = 2.187 pg Fe/cell), indicating reduced IONP associations, while the aggregates of VivoTrax and VivoTrax Plus likely distorted MPI data, resulting in elevated signals (VivoTrax = 48.52 pg Fe/cell; VivoTrax Plus = 220.3 pg Fe/cell) (Figure 3D).

**Figure 3.**
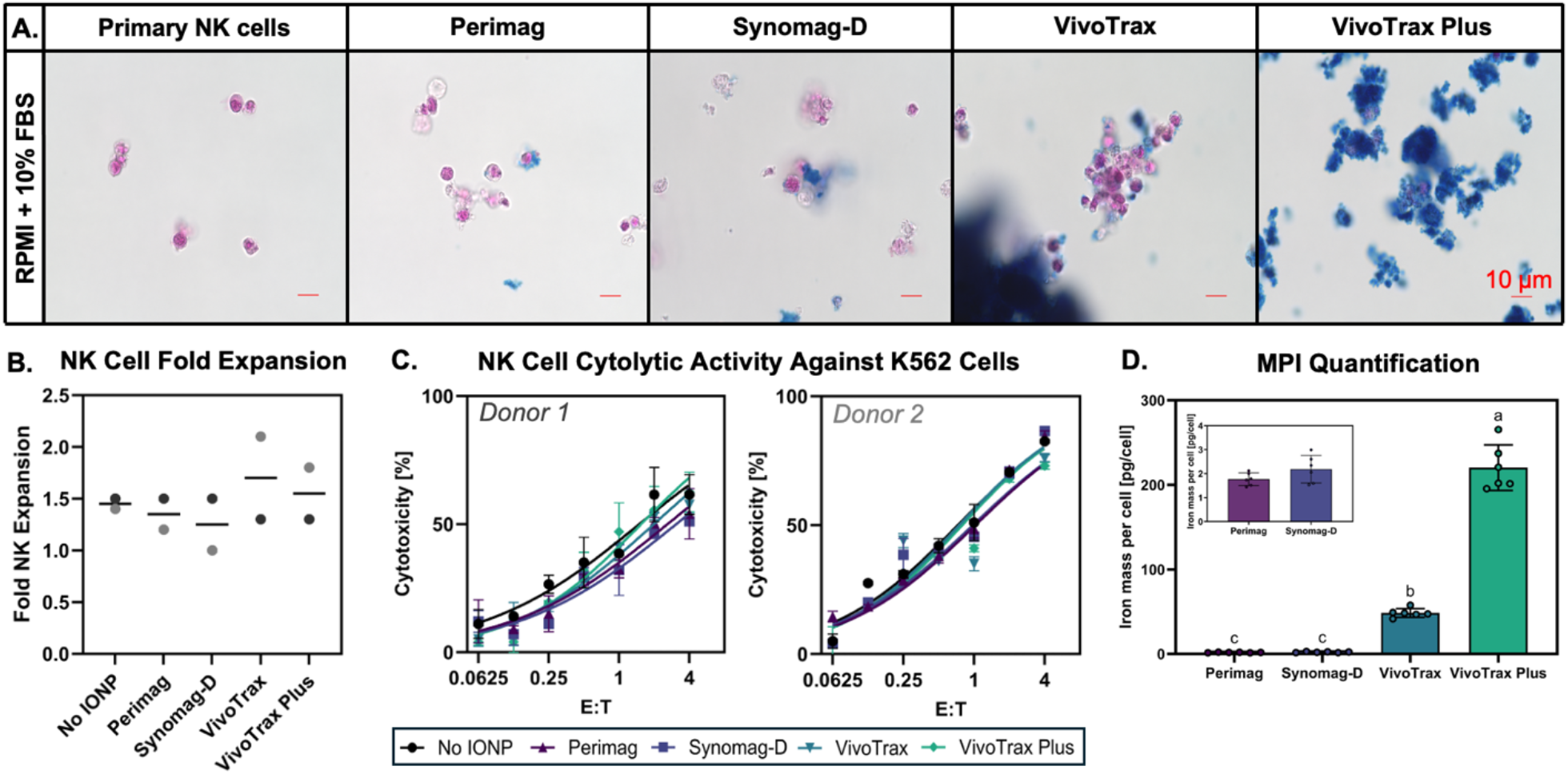
Iron Oxide Nanoparticle Interactions with Primary NK Cells (A) Prussian blue staining shows iron (blue) associated with primary NK cells (pink) after labeling with 100 μg Fe IONPs for 24 hours (scale bar = 10 μm) in RPMI 1640 supplemented with 10% FBS. (B) Fold expansion of NK cells from two different donors after incubation with 100 μg Fe/mL of IONP in various culture medias for 24 hours at 37°C and 5% CO_2_. (C) Multiple E:T ratios were used to determine single donor, dose-dependent cytotoxicity curves against K562 cells. (D) Iron content per cell, quantified by MPI. Different letters indicate statistically significant differences among groups (one-way ANOVA with a Tukey’s multiple comparison test, p < 0.05).

### Measurement of NK Cell Activity Against Solid Tumor Spheroids

NK cells efficiently recognize and lyse K562 cells due to the absence of MHC class I molecules on these targets, which enables robust activation through natural cytotoxicity receptors[41]. To further evaluate the functional integrity of IONP-labeled NK cells, cytotoxicity assays were extended to two non-small cell lung cancer (NSCLC) cell lines: A549 (harboring a KRAS mutation) and H1299 (a metastatic line derived from lymph nodes)[42]. These models were selected to mimic the immunosuppressive and heterogeneous environment of solid tumors, providing a more stringent test of NK cell cytolytic function.

Prior to co-culture experiments, we first assessed the direct cytotoxicity of IONPs on NSCLC spheroids in the absence of NK cells. Tumor spheroids were incubated overnight with 100 μg Fe/mL IONPs in serum-supplemented media. The next day, particles were serially diluted to match concentrations used during NK cell labeling and applied to GFP-expressing tumor spheroids. Spheroid growth and viability were monitored at 4-hour intervals using an IncuCyte SX3 live-cell imaging system. Among the tested formulations, VivoTrax and VivoTrax Plus exhibited high levels of apparent cytotoxicity, preventing further analysis in co-culture settings. This apparent toxicity may have resulted from nanoparticle aggregation interfering with GFP signal detection or from intrinsic cytotoxic effects of the particles themselves (Supplemental Figure S4).

NK cell-mediated cytotoxicity was assessed using NK cells pre-incubated with Perimag or Synomag-D, both of which demonstrated good colloidal stability and minimal direct apparent toxicity. Across both donors and in both NSCLC models, IONP-labeled NK cells retained their ability to kill tumor spheroids effectively. No statistically significant differences in cytotoxicity were observed when compared to unlabeled NK cells, indicating that Perimag and Synomag-D labeling does not impair NK cell effector function in the context of solid tumor targets (Figure 4).

**Figure 4.**
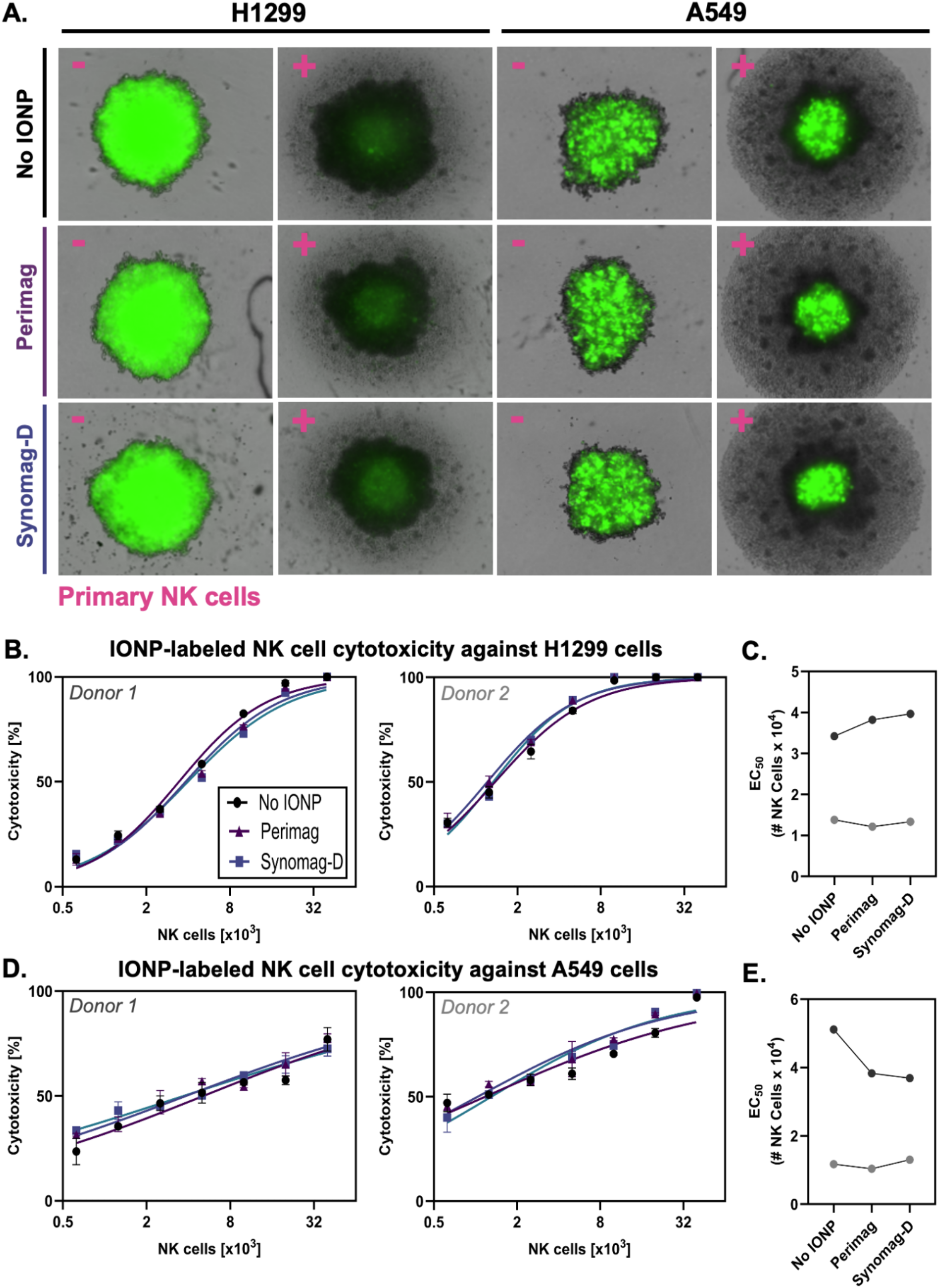
NK Cell Function Against Solid Tumor Spheroids (A) Representative IncuCyte images of spheroid cytotoxicity assay. GFP+ tumor spheroids were generated and cultured for 3 days prior to co-culture with IONP and/or NK cells. Pink plus/minus symbols indicate the presence or absence of 20,000 NK cells per condition. Images correspond to the 72 h (H1299) and 96 h (A549) time points. (B) NK cell cytotoxic activity against H1299 spheroids following incubation with 100 μg Fe/mL of Perimag and Synomag-D nanoparticles, compared to NK cell controls without nanoparticles. (C) Corresponding EC_50_ values calculated from the curve fit; lines connect matched donor conditions. (D) NK cell cytotoxic activity against A549 spheroids under the same nanoparticle treatment conditions. (E) EC_50_ values for A549 spheroid killing calculated from the curve fit; lines connect matched donor conditions. No statistical differences in cytotoxicity were determined by RM one-way ANOVA with a post-hoc Tukey’s test correction for multiple comparisons and by multiple paired t-tests for comparison of EC_50_ values.

### Removal of Contaminating Extracellular Iron Aggregates

To address signal overestimation caused by extracellular iron aggregates, a Ficoll-Paque density gradient was applied using VivoTrax Plus, the formulation with the greatest aggregation, as a proof-of-concept (Figure 5A and Supplemental Figure S5). The separation effectively removed extracellular iron from NK92MI cells and reduced large aggregates in primary NK cells, though small extracellular particles remained visible (Figure 5B). Approximately 85% cell loss was observed during the process (Figure 5C), limiting its applicability in high-yield workflows. Post-separation, MPI quantification showed reduced iron content per cell, confirming that extracellular iron contributes to signal inflation. Final labeling yielded 4.8 pg Fe/cell in NK92MI and 11.2 pg Fe/cell in primary NK cells, exceeding previously reported values for VivoTrax[31](Figure 5D).

**Figure 5.**
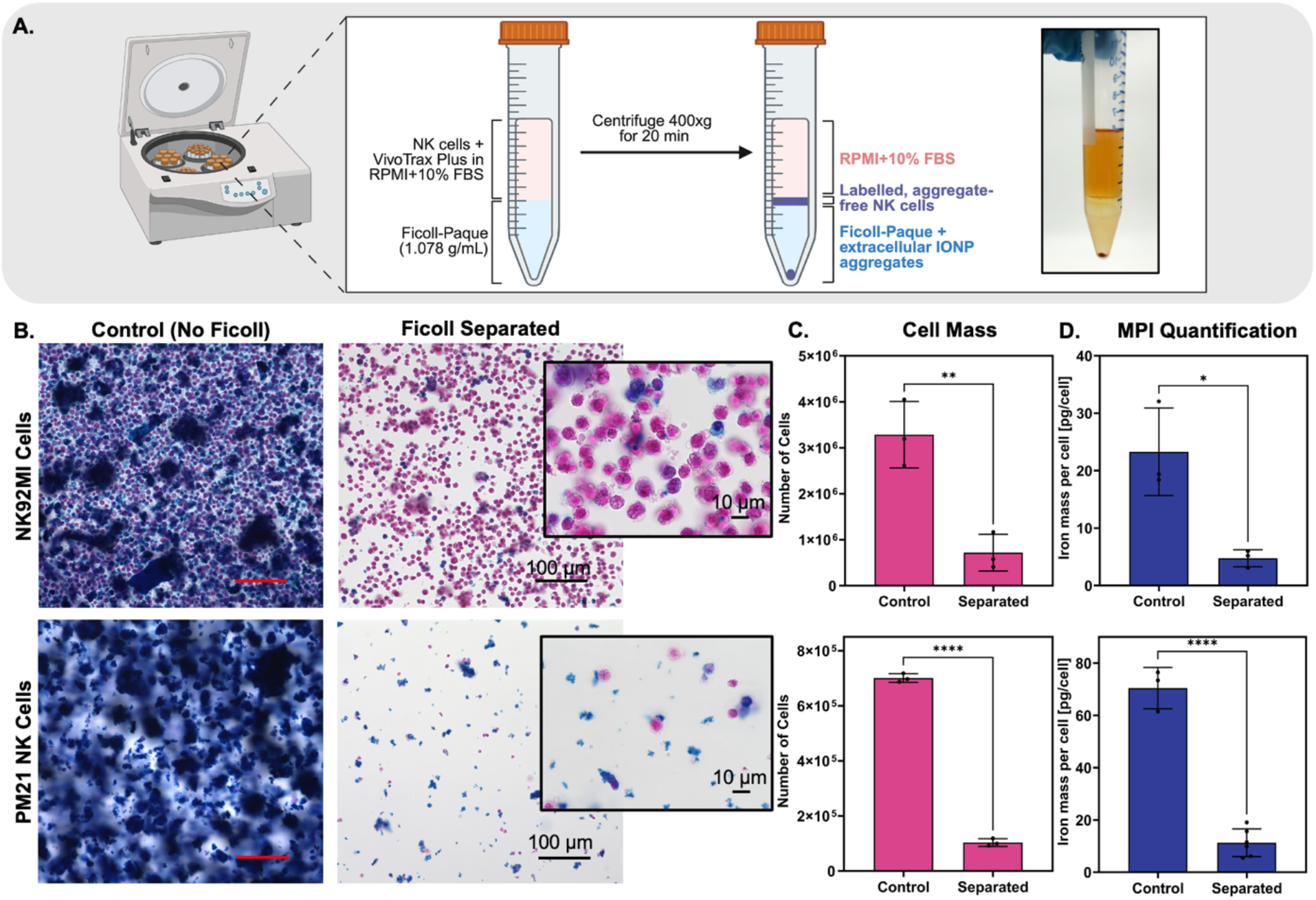
Density Gradient Separation to Remove IONP Aggregates. (A) Schematic detailing Ficoll-Paque density separation of iron aggregates from NK cells incubated with 100 μg Fe/mL of VivoTrax Plus in RPMI 1640 supplemented with 10% FBS. (B) Prussian blue staining shows the amount of extracellular iron (blue) aggregates and NK cells (pink) before and after Ficoll separation. Images were captured using the 20x objective (scale bar – 100 μm), inset shows images captured with the 100x objective (scale bar = 10 μm). (C) Number of cells and (D) iron content per cell, quantified by MPI, before and after Ficoll-Paque density separation. Statistical significance was determined using an unpaired Student’s t-test, where *p<0.05, **p<0.01, and ****p<0.0001.

### Balancing Labeling Efficiency with Cell Recovery for MPI Tracking

Assuming a detection threshold of 30 ng Fe for Synomag-D[43] and 34 ng Fe for VivoTrax Plus based on relative signal intensities, the minimum number of primary NK cells required for detection would be approximately 1.4×10^4^ cells for Synomag-D (at 2.2 pg Fe per cell) and 3×10^3^ cells for VivoTrax Plus (at 11.2 pg Fe per cell). However, after accounting for the 85% cell loss during Ficoll-Paque separation, which is used to remove extracellular aggregates from VivoTrax Plus samples, the required starting number of cells increases to approximately 2×10^4^, which exceeds the number of cells needed for Synomag-D-labeled cells. These calculations are summarized in Table 1.

**Table 1.** Estimated number of primary NK cells required for MPI detection, based on per-cell iron content and assumed limits of detection (LoD). Adjusted values for experiment reflect 85% cell loss during Ficoll-Paque separation.

| IONP | LOD (ng Fe) | Iron per Cell (pg Fe) | Cells Required for Detection | Cells Required for Experiment |
| --- | --- | --- | --- | --- |
| Synomag-D | 30 | 2.2 | $1.4 \times 10^4$ | $1.4 \times 10^4$ |
| VivoTrax Plus | 34 | 11.2 | $3 \times 10^3$ | $2 \times 10^4$ |

### IONP-labeled NK Cell Tracking in Anatomically Correct Phantoms

Imaging performance was assessed using an anatomically accurate mouse phantom with NK cells labeled with Synomag-D or density-separated VivoTrax Plus IONPs, distributed between the liver, which reflects the primary site of exogenous cell accumulation after systemic delivery, and a secondary region, which reflect a target site. This setup simulates in vivo conditions where liver accumulation may obscure target-site signal or reduce quantitative accuracy due to spillover. Target regions included the brain (anatomically distant from the liver) and the lung (in close proximity), the latter representing a more challenging scenario for detection and quantification[44].

To evaluate the impact of liver spillover on quantification, two segmentation strategies were applied to the brain and lung ROIs: a 0.5Max signal threshold (red contour in Figure 6) and a fixed 7-mm diameter circle centered at the local maximum (green contour in Figure 6). For each ROI, total signal was compared under two conditions: (1) capillary-only, with cells in the capillary and no liver signal, and (2) capillary + liver, with cells in the capillary located either in the brain or the liver. This approach isolates the influence of liver signal on the target region while keeping the ROI constant[40]. For each phantom, 5% of the total NK cells were placed in the capillary and 95% in the liver to mimic tumor homing, with the total cell number held constant across phantoms to compare nanoparticle performance in MPI.

**Figure 6.**
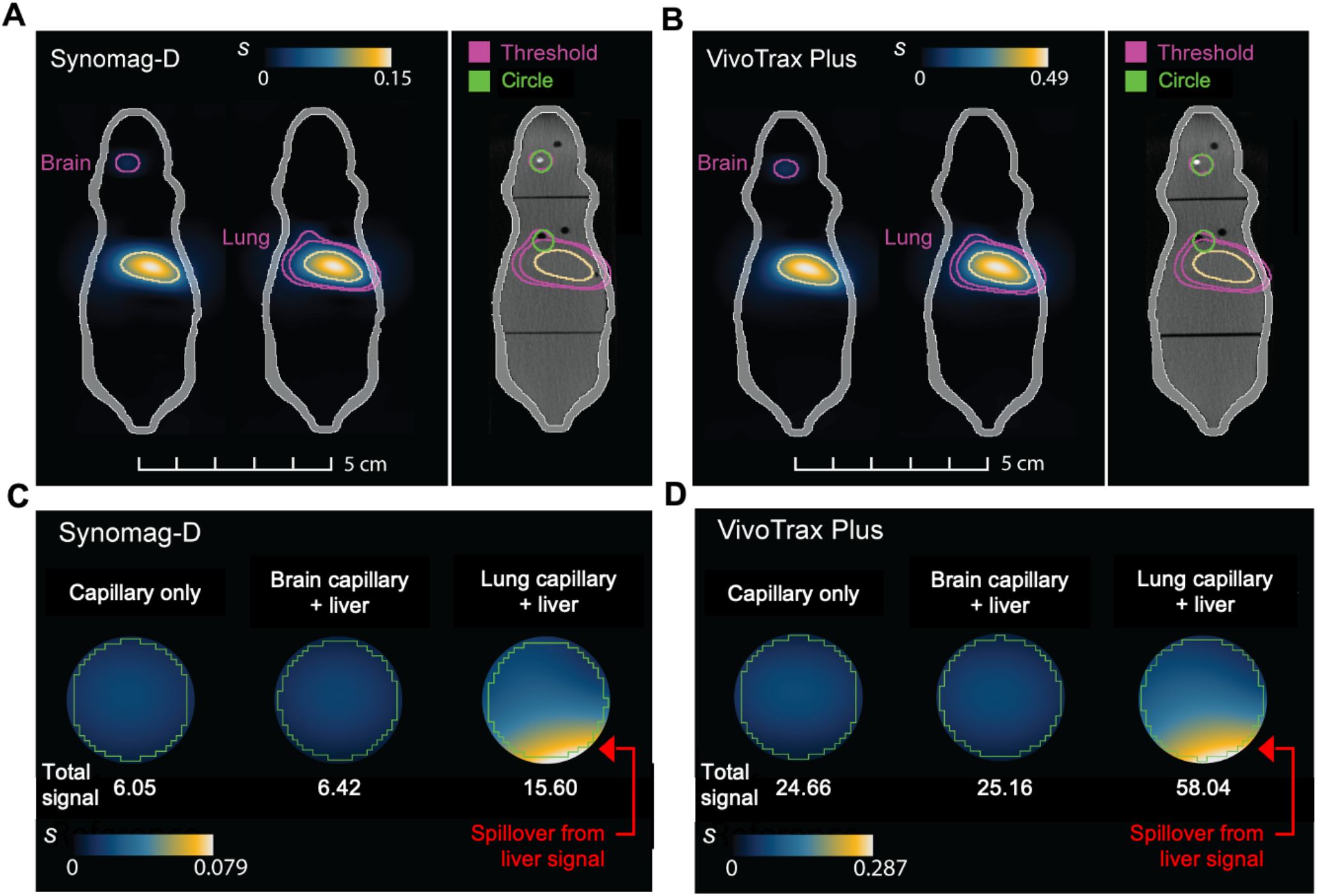
MPI scans of mouse phantoms with a liver cavity and a capillary positioned in the brain/lung region, both filled with NK cells labeled with (A) Synomag-D and (B) VivoTrax Plus (Ficoll-separated). Segmentations using a 0.5Max threshold or a fixed-area circle are overlayed on computed tomography images of the phantom for spatial reference. Total signal (arbitrary units) quantified within the fixed-area circle demonstrates artificial signal inflation from the liver in the lung region for phantoms containing (C) Synomag-D and (D) VivoTrax Plus (Ficoll-separated) tracers.

Segmentation of the brain signal was straightforward under both conditions. In contrast, although the lung signal was visually identifiable, threshold-based segmentation could not be applied without including spurious signal from the adjacent liver (Figure 6). The distorted threshold shape highlights substantial liver signal spillover, emphasizing the effect of distance from the liver on MPI-based cell tracking. Fixed-area spherical segmentation provided a more robust alternative by limiting spillover and enabling consistent quantification (Figure 6 and Supplemental Figure S6).

Quantification confirmed these observations. In the brain, liver signal had minimal impact: for Synomag-D-labeled NK cells, total signal in the capillary + liver condition was 1.02x and 1.06x that of the capillary-only condition using thresholded and fixed-area segmentations, respectively, while VivoTrax Plus-labeled cells showed signal that was 1.02x that of the capillary-only condition for both methods. Ratios close to 1 indicate negligible liver-induced bias. By contrast, lung-region quantification was strongly affected by liver spillover (Figure 6C and 6D). Using fixed-volume segmentation, the total signal in the lung ROI was overestimated, reaching 2.57x and 2.35x that of the capillary-only condition for Synomag-D- and VivoTrax Plus-labeled NK cells, respectively, demonstrating substantial spillover from the liver.

When comparing tracer performance more generally, VivoTrax Plus-labeled NK cells (after Ficoll separation) contained approximately 5x more iron per cell than Synomag-D-labeled cells. Accordingly, the total MPI signal from VivoTrax Plus cells was ∼3.8x higher than that from Synomag-D-labeled cells, independent of segmentation method and consistent across both capillary-only and liver-only conditions (Figure 6A and 6B). This confirms that higher per-cell iron content yields a stronger signal in mouse phantoms, consistent with linear scaling of MPI signal, though deviations may occur in a cellular context.

## Discussion

Tracking NK cells non-invasively remains a major unmet need in the field of cancer immunotherapy. While NK cell therapies are gaining traction due to their unique advantages in targeting tumors without causing graft-versus-host disease, their in vivo migration and persistence are not well understood[8]. Sehl et al. reported the use of MPI to track NK cells, demonstrating the feasibility of labeling NK cells with an MPI tracer and imaging of locally administered cells in cadavers but leaving open questions about optimal labeling methods and particle selection[31]. In this study, we expanded on this foundation by labeling NK92MI cells and, for the first time, primary human NK cells, using commercially available IONPs for MPI tracking. We confirmed that labeled NK cells were viable and maintained functional cytotoxicity, including activity against challenging three-dimensional tumor spheroids. Importantly, both cell types were detectable by MPI, validating the potential of this imaging modality for real-time, non-invasive tracking of adoptively transferred NK cells.

A key finding of our work is the complex interplay among particle colloidal stability, particle-cell association, and the need for post-labeling purification. While VivoTrax and VivoTrax Plus particles offered stronger MPI signals due to higher iron associations, their tendency to aggregate extracellularly necessitated Ficoll-Paque density gradient separation to remove free iron. This purification step led to a significant loss of viable NK cells, reducing the overall cell yield and offsetting the initial signal advantage. In contrast, Perimag and Synomag-D particles demonstrated superior colloidal stability with minimal aggregation, simplifying the labeling workflow and preserving more cells, albeit at a lower per-cell MPI signal intensity. It should be noted that this feature is not unique to MPI but has also been observed in other imaging modalities, including MRI[18]. Therefore, the choice of IONP formulation should reflect the specific demands of the study, balancing labeling efficiency, signal strength, and processing constraints.

Mouse phantom studies demonstrate that these differences extend beyond labeling efficiency and influence quantitative imaging. They also provide a practical, animal-free platform to study imaging and cell-tracking parameters under controlled conditions[40]. In this model, liver signal spillover led to pronounced overestimation in nearby regions such as the lung, while distant regions like the brain remained largely unaffected and reliably quantifiable. This is not an MPI-specific limitation but a fundamental challenge in preclinical cell tracking: accumulation in clearance organs such as the liver can dominate signal and compromise accurate localization and quantification at target sites[44–46]. The issue is especially pronounced in small animal models, where organ proximity exacerbates spillover. However, it is not limited to mice and may impact larger animals and clinical settings, especially for targets located near the liver. Overall, these results underscore the need to account for spatial context and spillover effects when interpreting imaging data and highlight that improving tracer design alone is insufficient without parallel advances in analysis strategies.

From a safety perspective, iron-based cell tracking using clinically approved particles is promising. The use of commercially available particles manufactured under Good Manufacturing Practice conditions supports regulatory compliance and patient safety[20]. Given the biocompatibility of these formulations and their prior use in human imaging, we anticipate that MPI-based NK cell tracking can be safely integrated into future clinical studies[47].

From a translational standpoint, Synomag-D may be better suited for large-scale or clinical use, where workflow simplicity, particle stability, and high cell recovery are prioritized. Its performance highlights how stable labeling, despite lower signal intensity, can result in more practical and reproducible imaging outcomes. On the other hand, VivoTrax Plus may retain value in studies where only small populations of cells migrate to target tissues, such as in the context of early solid tumor infiltration. Its higher per-cell iron content allows for more sensitive detection of rare cell events, despite added purification requirements.

A notable limitation of this study is that evaluations were confined to short-term in vitro experiments. Future work will need to explore longitudinal tracking of NK cells in vivo using tumor-bearing mouse models to better understand the relationship between MPI signal intensity, NK cell localization, and therapeutic outcomes. It is important to recognize that IONP-based tracking inherently allows only short-term monitoring due to signal dilution from cell proliferation and natural nanoparticle degradation[11,15]. However, given that many adoptive NK cell therapies currently exhibit limited in vivo persistence and proliferation, this short-term tracking may still provide valuable insights[48]. Despite these constraints, IONP-based MPI remains a promising tool for capturing the critical early phases of NK cell biodistribution and tumor targeting.

More broadly, these results suggest that current commercial IONPs are not yet optimized for NK cell labeling. Future nanoparticle designs should aim to enhance intracellular uptake specifically in NK cells, possibly through surface modifications that facilitate membrane fusion or receptor-mediated internalization. Strategies such as incorporating cell-penetrating peptides or NK-targeted ligands could improve labeling efficiency without inducing cytotoxicity. At the same time, formulations must maintain high magnetic performance and robust colloidal stability in serum-containing environments. These design considerations will be especially important as MPI technology continues to evolve toward human-scale scanners[24,26] and potential clinical applications.

## Conclusion

In summary, this work provides a critical evaluation of commercially available IONPs for magnetic particle imaging of NK cells, marking the first reported labeling of primary human NK cells for MPI tracking. By comparing Synomag-D and VivoTrax Plus, we reveal key tradeoffs between nanoparticle colloidal stability and per-cell iron associations, which directly impact detection sensitivity and workflow practicality. Complementary mouse phantom studies further demonstrate that these differences extend to imaging performance in biologically relevant conditions, where liver signal spillover can significantly bias quantification in nearby target regions. These insights inform the selection of labeling strategies that balance signal intensity, cell viability, and processing complexity, addressing a significant gap in non-invasive NK cell tracking technology. Ultimately, our findings lay the groundwork for refining MPI-based approaches that can accurately monitor NK cell biodistribution and persistence in vivo, thereby advancing both mechanistic understanding and clinical translation of NK cell immunotherapies.

## Acknowledgements

This work was supported by the National Science Foundation (NSF) Division of Chemical, Bioengineering, Environmental, and Transport Systems (CBET) Faculty Early Career Development (CAREER) Program (Award 1845728), Graduate Research Fellowship Program (Award 1000308740), and the Leo Clair and Robert Adenbaum Foundation. The content is solely the responsibility of the authors and does not necessarily represent the official views of the National Science Foundation. Research in the Rinaldi-Ramos lab was supported by the National Institute for Biomedical Imaging and Bioengineering (NIBIB) (R01EB031224) and the National Cancer Institute (NCI) (R01CA298804). Research in the Copik lab was supported by FL DOH James and Ester King Program (Grant No. 25JK07 and 9JK04). Figure 5A was created in Biorender.com under a premium license. The authors also thank all laboratory members who contributed to the initial studies underlying this publication.

