## Supplemental for "Towards Reliable Tracking of Natural Killer Cells Using Commercial Iron Oxide Nanoparticles and Magnetic Particle Imaging"

### ABSTRACT

Non-invasive tracking of natural killer (NK) cells remains a major challenge in cancer immunotherapy, limiting our understanding of their in vivo migration and persistence. Magnetic particle imaging (MPI) offers a quantitative, real-time method for visualizing labeled cells, yet optimal labeling protocols for NK cells have not been established. Here, we evaluate commercially available iron oxide nanoparticles (IONPs) for MPI labeling of both NK92MI cells and primary human NK cells. Labeled cells retained viability and cytotoxicity, including activity against three-dimensional tumor spheroids, and were detectable by MPI. To further examine imaging performance in a biologically relevant context, we employed mouse phantoms that recapitulate organ-specific signal distributions, enabling evaluation of quantification and liver spillover effects. We identify key tradeoffs between particle colloidal stability and per-cell iron content: VivoTrax and VivoTrax Plus provided higher MPI signal but required post-labeling purification, reducing cell recovery, whereas Synomag-D and Perimag were more stable and preserved cell yield despite lower signal intensity per cell. These results provide a framework for selecting nanoparticles that balance detection sensitivity, cell viability, and workflow practicality, advancing non-invasive NK cell tracking.

| | |
|---|---|
| Supplemental Figure 1: Standard Curves Correlating MPI Signal to Know Iron Masses | Page 2 |
| Supplemental Figure 2: IONPs in Various NK Cell Culture Media | Page 3 |
| Supplemental Figure 3: Prussian Blue Staining at 4 hours | Page 4 |
| Supplemental Figure 4: Toxicity of IONPs on NSCLC Spheroids | Page 5 |
| Supplemental Figure 5: Pilot Ficoll Separation Study | Page 5 |
| Supplemental Figure 6: Quantification of MPI Signal from Mouse Phantoms | Page 6 |

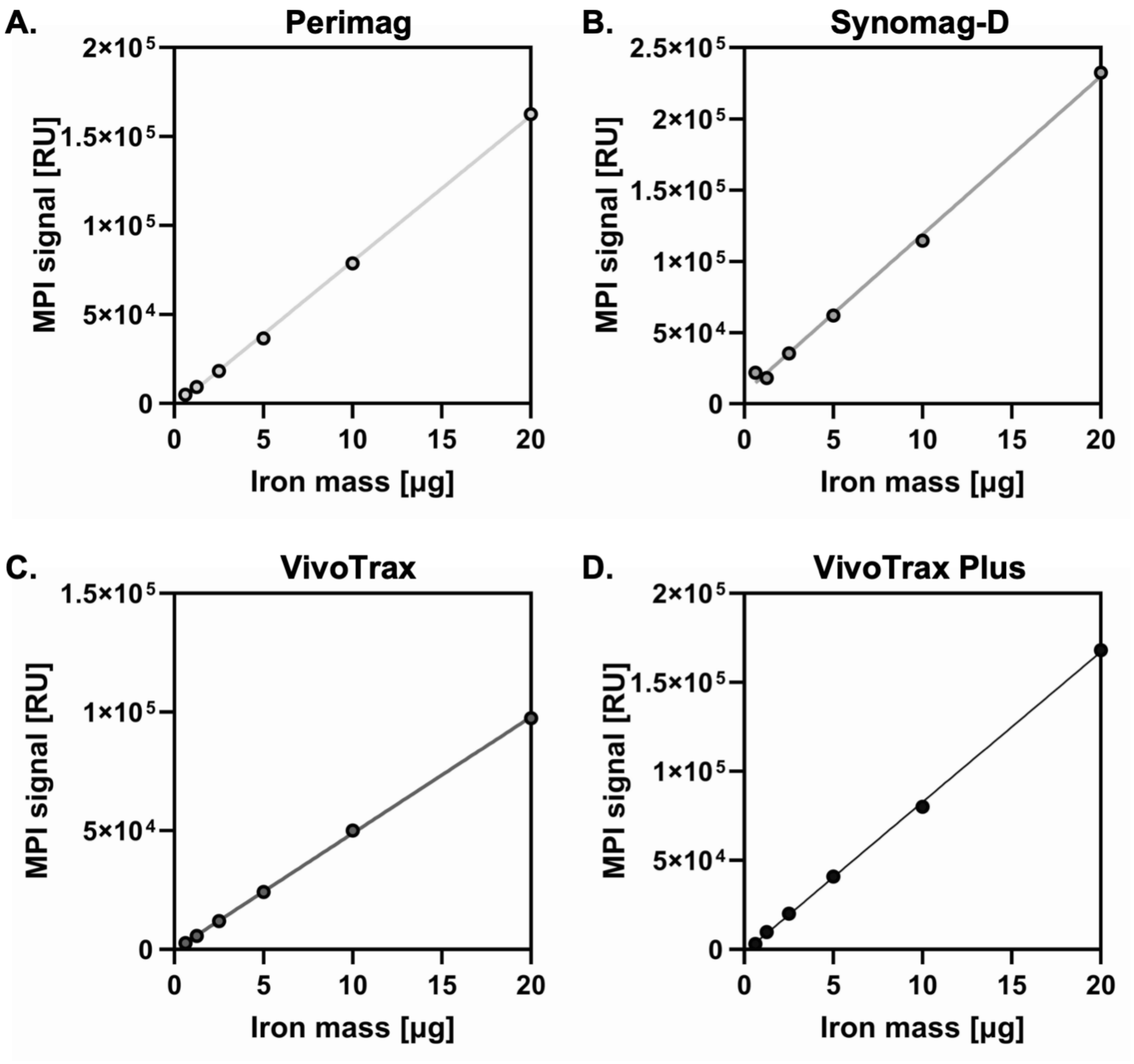


**Supplemental Figure 1.** PI signal was measured for serial dilutions of IONPs, and linear regression was used to generate standard curves for: (A) Perimag, (B) Synomag-D, (C) VivoTrax, and (D) VivoTrax Plus. These calibration curves were used to quantify intracellular iron content in labeled NK cells via MPI.

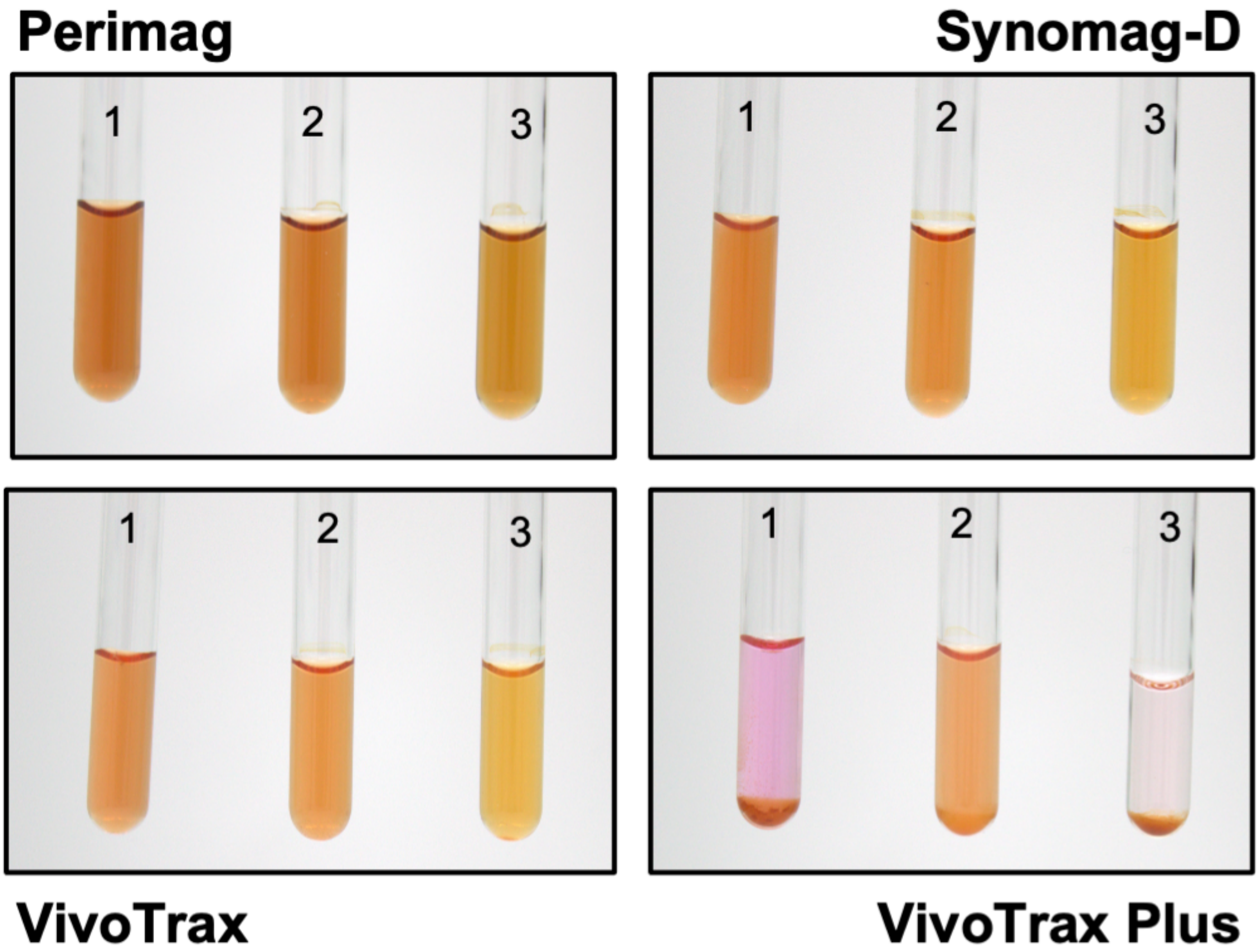


**Supplemental Figure 2.** Commercial IONPs in various NK cell culture media. Conditions 1, 2, and 3 correspond to: (1) RPMI alone, (2) RPMI supplemented with 10% fetal bovine serum (FBS), and (3) OptiMEM supplemented with 1% ITS.

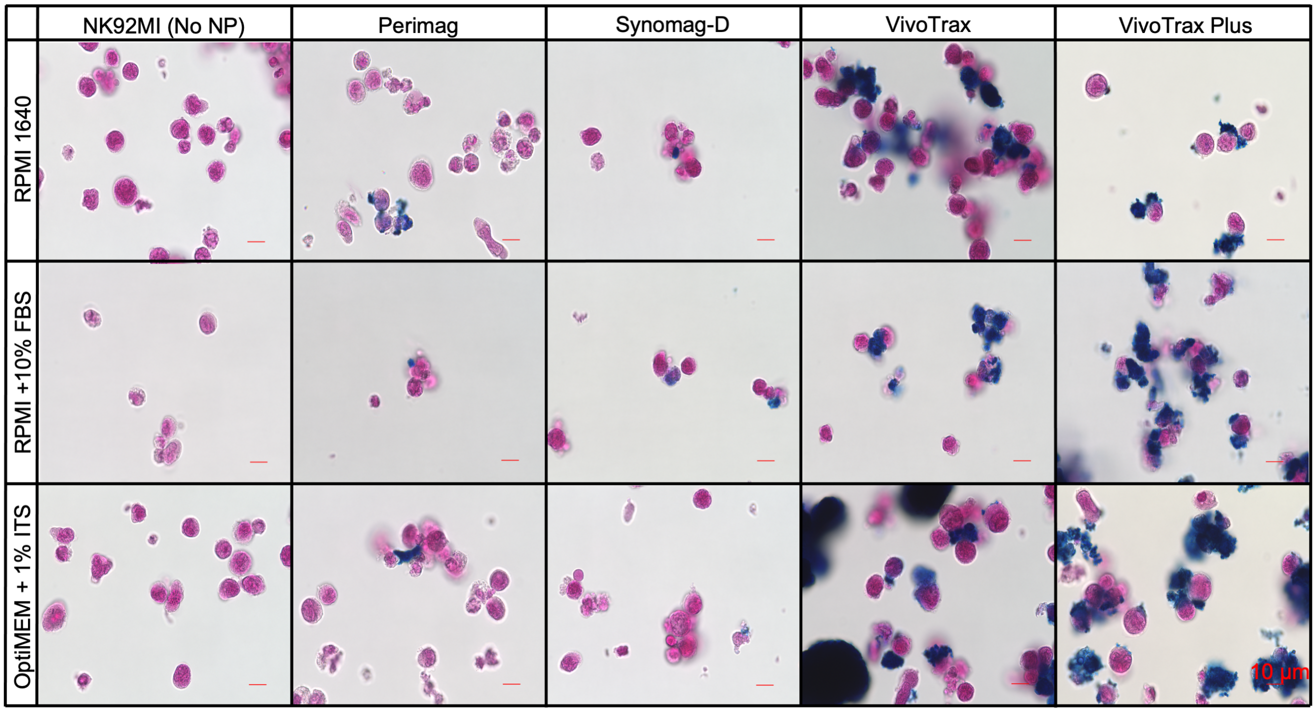


**Supplemental Figure 3.** Prussian blue staining of IONP (100 $\mu$g Fe/mL) interactions with NK92MI cells after 4 hours of co-culture (scale bar = 10 $\mu$m).

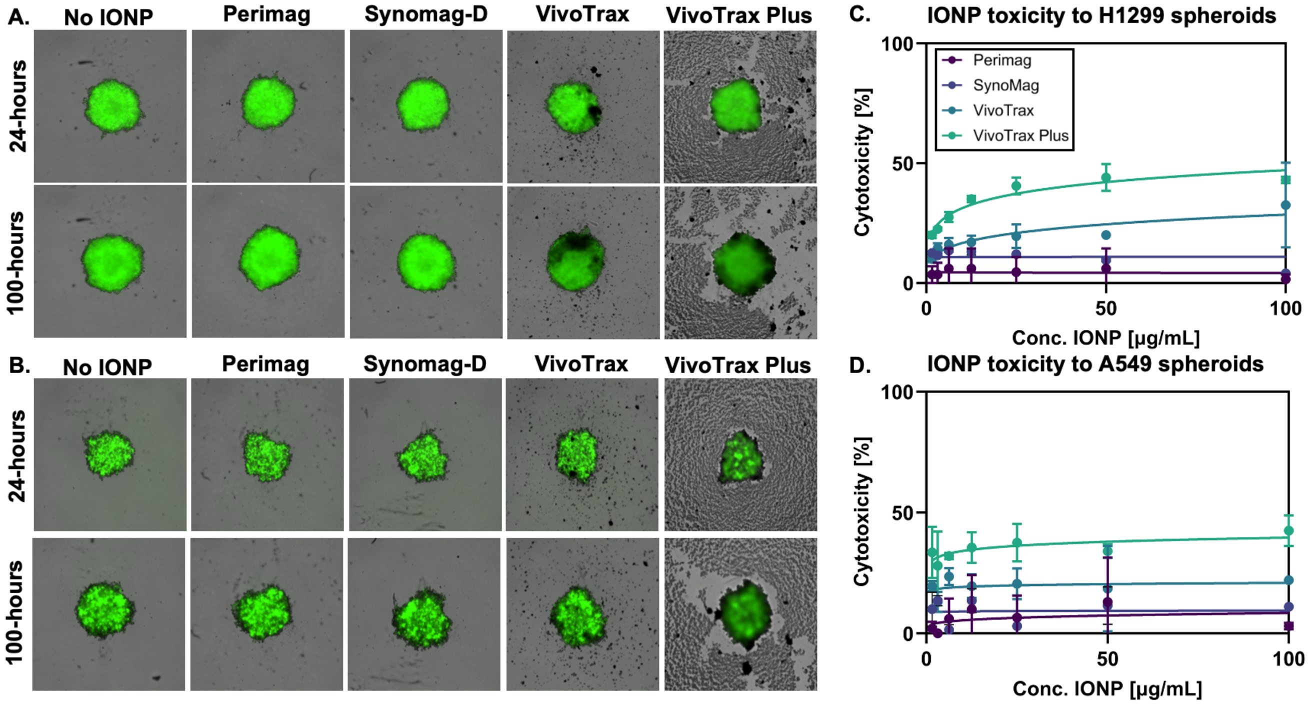


**Supplemental Figure 4.** Toxicity of IONPs on NSCLC spheroids in the absence of NK cells. IONPs were incubated overnight in serum-containing media at 100 $\mu$g Fe/mL. The following day, particles were diluted using the same protocol applied for NK cell labeling, and added to GFP-expressing NSCLC spheroids at the indicated concentrations. Representative images of tumor spheroids were acquired every 4 hours using an IncuCyte SX3 live-cell imaging system for (A) H1299-GFP+ and (B) A549-GFP+ spheroids. (C,D) Quantification of spheroid size and GFP expression demonstrates IONP-induced cytotoxicity of VivoTrax and VivoTrax Plus in both H1299 and A549 models.

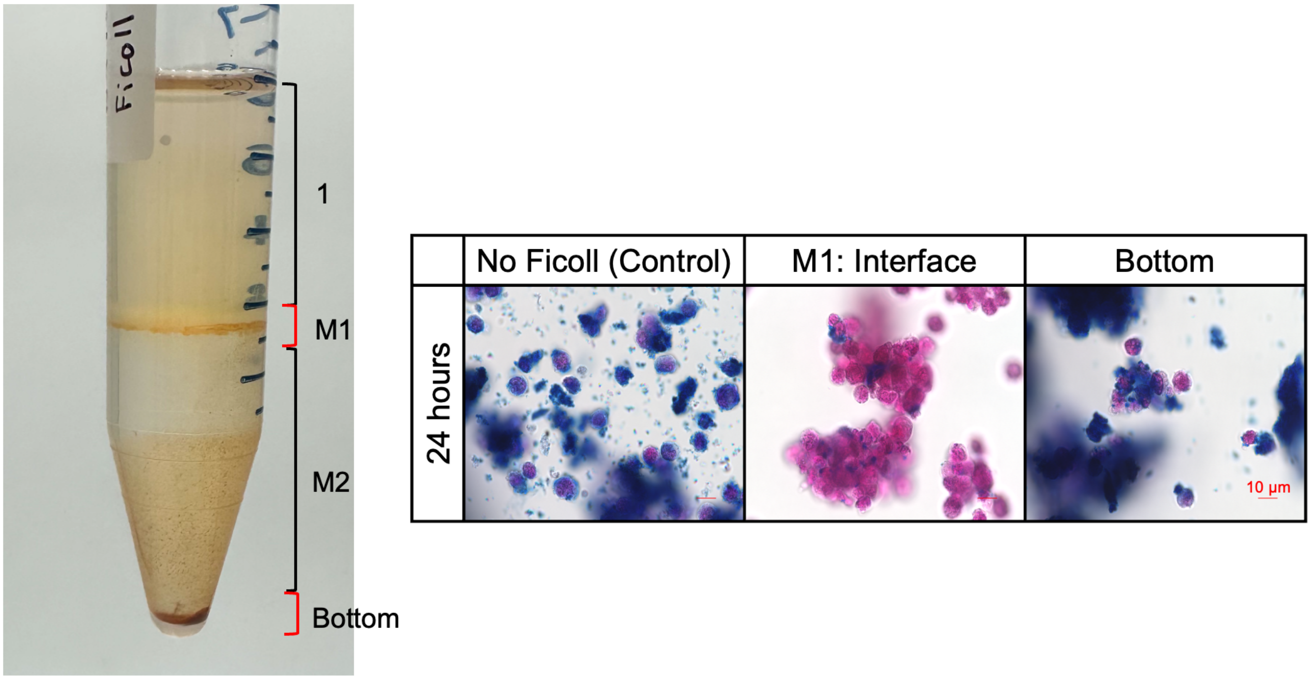


**Supplemental Figure 5.** Pilot Ficoll separation study. NK92MI cells were cultured with 100 $\mu$g Fe/mL VivoTrax Plus in OptiMEM supplemented with 1% ITS for 24 h. Samples collected from different regions of the interface were stained with Prussian blue and nuclear fast red to visually assess iron uptake and purification efficiency (scale bar = 10 $\mu$m).

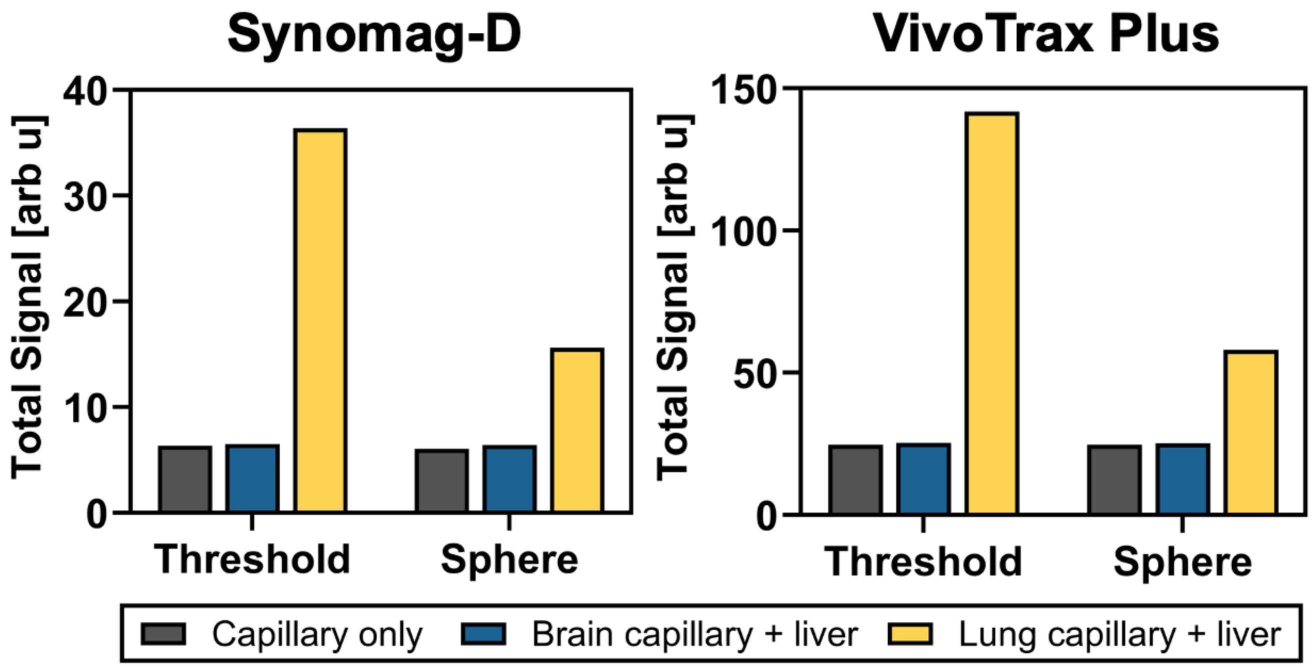


**Supplemental Figure 6.** Quantification of MPI signal from NK cells labeled with Synomag-D and VivoTrax Plus (Ficoll-separated) tracers in anatomically correct phantoms using different thresholding methods. To evaluate the impact of liver spillover on quantification, two segmentation strategies were applied to the brain and lung ROIs: a 0.5Max signal threshold (red contour in Figure 6) and a fixed 7 mm3 sphere centered at the local maximum (green contour in Figure 6). For each ROI, total signal was compared under two conditions: (1) capillary-only, with cells in the capillary and no liver signal, and (2) capillary + liver, with cells in the capillary located either in the brain or the liver.
